# Expression patterns of Arc mRNA after renewal of appetitive behavior in female rats

**DOI:** 10.1101/2021.07.20.453088

**Authors:** Emily N Hilz, Laura A. Agee, Donyun Jun, Marie H Monfils, Hongjoo J. Lee

## Abstract

Renewal of appetitive behavior depends on the gonadal hormonal state of the female rat. In this experiment the effect of female rat estrous cycle stage on renewal of appetitive behaviors is replicated and extended upon to understand how endogenous hormonal states around the estrous cycle drive renewal at the neuronal population level. Estrous cycle stage (i.e., proestrus (P, high hormone) or metestrus/diestrus (M/D, low hormone)) was considered during two important learning and behavioral expression windows: at extinction training and during LTM/renewal testing. First, rats in P during context-dependent extinction training but in some other stage of the estrous cycle during long-term memory and renewal testing (Different) were shown to exhibit more renewal of conditioned foodcup (but not conditioned orienting) behavior compared to rats in other estrous cycle groups. Next, cellular compartment analysis of temporal activity using fluorescence *in situ* hybridization (catFISH) was used to examine immediate early gene activity of *Arc* mRNA in neuronal populations after distinct context-stimulus exposures (i.e., extinction and acquisition test contexts). *Arc* mRNA expression patterns were examined in the prefrontal cortex (PFC), amygdala, hippocampus (HPC), and paraventricular nucleus of the thalamus. P-different rats showed differential neuronal population activity in the infralimbic cortex of the PFC, the lateral amygdaloid nucleus, and both CA1 and CA3 regions of the dorsal HPC. In each region P-different rats exhibited more co-expression and less specificity of *Arc* mRNA compared to other hormonal groups, indicating that renewal of appetitive foodcup behavior induces *Arc* mRNA in overlapping neuronal populations in female rats.

## Introduction

Extinction can be used to reduce the occurrence of maladaptive behaviors; while useful as a treatment tool, the generalization of extinction training from the rehabilitative context to the home context can present a therapeutic challenge. Maladaptive behaviors are learned in specific contextual environments, and these environmental cues can serve as a prompt for reward availability. Oftentimes extinction training occurs in an environment separate from the one in which the initial conditioned stimulus-unconditioned stimulus (CS-US) association was formed; rehabilitative failure can be attributed in part to the differences between the rehabilitative context and the home environment. Cues that are extinguished within the rehabilitative context can regain their reward-associative power when presented in the original acquisition context (Wing and Shoaib, 2008; Bouton and Peck, 1989), leading to resurgence of behavior in a phenomenon known as renewal. Renewal is the resurgence of extinguished behavior dependent on context stimuli.

Female rats exhibit less renewal compared to males; moreover, renewal of appetitive behavior in female rats is influenced by female sex steroid hormones in ovariectomy (OVX) models (wherein exogenous estradiol treatment enhances renewal of food-seeking behavior; Anderson and Petrovich, 2017; 2015). We have extended this to show that female renewal is influenced by endogenous sex steroid hormonal states associated with stages of the rat estrous cycle. In rats, the estrous cycle is a four- or five-day period over which female hormones fluctuate throughout the brain and body. Each independent stage of the cycle is associated with specific gonadal hormone levels (for review see Butcher et al., 1974) and can be identified by the presence of certain cell structures and concentrations in the vaginal epithelium (Marcondes et al., 2002). Rats in proestrus (P, high hormonal stage) during extinction training, but some other low-hormone stage of the estrous cycle during renewal testing (Different), exhibited enhanced renewal compared to rats trained and tested in other hormonal stages (Hilz et al., 2019). Cellular functional activity after renewal in the hippocampus (HPC), paraventricular nucleus of the thalamus (PVT), and amygdala was enhanced among these animals.

The purpose of the current experiment is to replicate and expand upon this previous finding to understand how estrous cycle stage modulates renewal of appetitive behavior. While the circuitry underlying the contextual modulation of conditioned responding is well established in both fear and appetitive conditioning, less is known about how neuronal ensembles in the HPC, PFC, and amygdala may guide the expression of previously extinguished behaviors. *Arc* catFISH (cellular compartment analysis of temporal activity using fluorescence *in situ* hybridization) allows examination of neuronal populations selectively activated in response to two temporally distinct events. *Arc* mRNA is an immediate early gene expressed after cell activation in either the cell nucleus and/or cytoplasm (Lee et al., 2016; Vazdarjanova et al., 2002; Guzowski et al., 1999); *Arc* mRNA migrates in a temporally predictable pattern from the cell nucleus to the cytoplasm over an established window of time (∼20 minutes), and the pattern of *Arc* mRNA expression can be used to identify unique and overlapping cell population activity in response to conditioned stimuli. Renewal testing induces context-dependent patterns of selective *Arc* mRNA expression in the amygdala and prefrontal cortex (PFC) of rats that have undergone context-dependent extinction training, while rats without context-dependent extinction training show proportionately greater numbers of non-selective neurons co-expressing both nuclear and cytoplasmic *Arc* mRNA (Orsini et al., 2013).

In this experiment the selectivity of *Arc* mRNA expression using catFISH after estrous-cycle dependent renewal is examined. Female rats underwent appetitive Pavlovian conditioning, context-specific extinction training, and then were tested for long-term memory (LTM) of extinction training and renewal of appetitive behavior in quick succession (20 minutes). Stages of the estrous cycle were considered during extinction training and at testing. Rats were then sacrificed and collected tissues were processed using catFISH for *Arc* mRNA. We hypothesized that estrous-cycle modulation of renewal in females may occur via better context representation in neuronal populations of the HPC or indicate a failure to retrieve extinction learning in a state-dependent manner via population activity in the PFC and/or amygdala. Characterizing the selectivity of *Arc* mRNA expression in these regions of the brain known to support renewal will help to identify how they regions respond to and represent different learning contexts and may indicate a potential mechanism of renewal in females.

## Methods

### Subjects

Thirty-four female Sprague-Dawley rats (Envigo; Indianapolis, IN, USA) weighing 200-275g were used in this study. Rats were pair-housed on a reverse 14:10 light:dark cycle with lights off at 10AM. Rats had *ad libitum* access to food and water for approximately one week after receipt; once acclimated to the colony room, rats received daily vaginal lavage and were food-restricted to reach approximately 90% free-feeding bodyweight for the entirety of the experiment. All procedures were conducted under the approval of the Institutional Animal Care and Use Committee at the University of Texas at Austin and in accordance with NIH guidelines.

### Apparatus

All procedures were conducted using acrylic-aluminum conditioning chambers measuring 30.5cm W / 25.5cm L / 30.5cm H (Coulbourn Instruments, Whitehall, PA). The chambers had clear acrylic front and back walls, aluminum siding and roof, and steel-rod flooring 0.5cm in diameter and spaced 1.0cm apart above a plastic drop-tray. A food-cup apparatus was located on the right side of the chamber 2.5cm above the floor and connected to an external magazine which delivered food-pellets (45mg TestDiet, Richmond, IN). Entries into the food-cup were measured automatically via breaks in an infrared beam across the lip of the food-cup. A 2-watt bulb was located 20cm above the floor on the right side of the chamber; illumination of this bulb served as the conditioned stimulus (CS) which predicted food-pellet delivery. Each chamber was enclosed in a sound- and light-attenuating box (Coulbourn Instruments, Whitehall, PA) which contained a wall-mounted camera used to record conditioning sessions.

### Behavioral Procedure

#### Estrous cycle determinations

The estrous cycle stage of each rat was monitored prior to and during experimental procedures using cytological examination of cells collected from the vaginal epithelium via vaginal lavage (40µl 0.9% saline flush). Samples were collected at the start of each dark cycle and immediately examined under 10x magnification prior to experimental procedures. Each stage of the estrous cycle is associated with specific gonadal hormone levels and can be identified by differing cell structures and concentrations (for review, see Marcondes et al., 2002). In short, proestrus is the stage of the estrous cycle associated with peaks in gonadal hormones such as estradiol and progesterone and is identifiable by a high concentration of nucleated epithelial cells. Metestrus and diestrus (sometimes referred to as diestrus 1 and diestrus 2) are transitional stages of the estrous cycle associated with generally low levels of gonadal hormones and characterized by a mix of cornified epithelial cells and leukocytes. Estrus, which was excluded from this experiment, is a more hormonally volatile stage of the estrous cycle during which hormone levels begin high and transition to low over the course of the stage. Estrus is identifiable by a high concentration of cornified epithelial cells.

#### Renewal Procedure

Behavioral procedures like those in Hilz et al., 2019 were used with slight modifications to allow for the utilization of FISH for *Arc* mRNA (Fig. 1). In short, rats underwent renewal procedures consisting of four appetitive conditioning sessions in context A, one extinction training session in context B, and tests for long-term memory (LTM) of extinction and renewal of appetitive behavior on the same day separated by 20 minutes (counterbalanced) in contexts B and A respectively. The counterbalanced order in which procedures were conducted was ABBA (i.e., acquisition, extinction, LTM, then renewal) or ABAB (i.e., acquisition, extinction, renewal, then LTM). Context A consisted of clear acrylic/aluminum walls, a steel-rod floor, and a neutral scent. Context B consisted of black and white horizontal-lined paper fastened outside the clear acrylic walls, a smooth black floor insert, and clean bedding identical to that used in the home-cage lining the drop-tray beneath the floor to introduce a non-neutral scent.

**Fig. 1.**
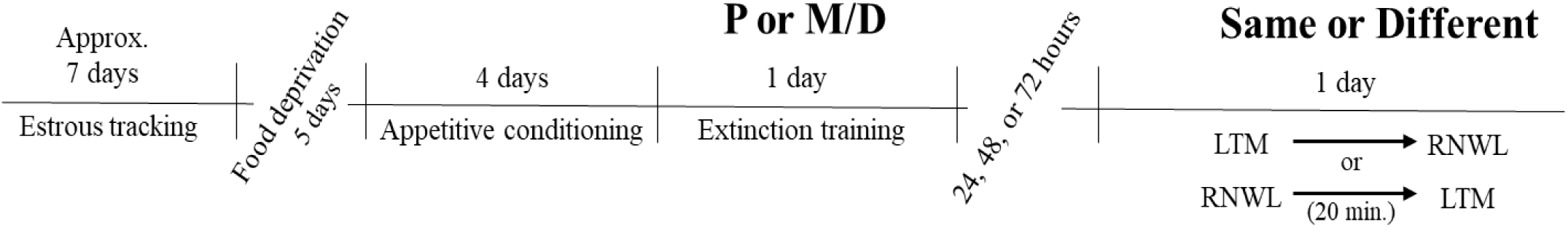
Timeline of behavioral manipulation used in renewal procedures. Renewal (RNWL) and long-term memory for extinction (LTM) tests are counterbalanced and separated by 20 minutes.

One day prior to appetitive conditioning rats were trained to retrieve and consume 30 food-pellets delivered on a 60s fixed interval schedule from the food-cup over a 30-minute session. Except for day 1, each appetitive conditioning session consisted of 16 10s light CS presentations paired with the delivery of a single noncontingent food-pellet over approximately 35 minutes on a 120s +/- 60s variable interval schedule. The first day of conditioning consisted of 8 consecutive unpaired light CS presentations (no food-pellet) and 8 consecutive light CS presentations paired with the delivery of a single food pellet on the same variable interval schedule. After appetitive conditioning rats underwent one extinction training session consisting of 18 unpaired CS presentations over approximately 40 minutes on a 120s +/- 60s variable interval schedule. Extinction training typically occurred two days after the final appetitive conditioning session to allow rats to cycle into the appropriate estrous cycle stage. Rats were in either proestrus (P, high hormone) or metestrus/diestrus (M/D, low hormone) during extinction training procedures. Rats in estrus were excluded from analysis due to the fluctuating nature of gonadal hormones during this stage of the estrous cycle. Finally, rats were tested for LTM of extinction and renewal of appetitive behavior in two separate testing sessions separated by 20 minutes on the same day. Each test consisted of 3 unpaired light CS presentations on a 120s +/- 60s variable interval schedule lasting approximately 7 minutes in contexts A and B. The order these tests were presented was counterbalanced. Rats were categorized as either “same” (i.e., in the same estrous cycle stage as extinction) or “different” (i.e., in a different estrous cycle stage from extinction) during testing. This produced four experimental groups: P-same, P-different, M/D-same, and M/D-different. Additionally, four home-cage control rats in either P or M/D (*n =* 2 per cycle stage, 4 total) with no LTM or renewal testing were sacrificed for baseline FISH comparison.

#### Scoring conditioned behavior

Appetitive conditioning was assessed using methods identical to Hilz et al., 2019. In short, foodcup behavior was measured automatically via breaks in an infrared beam at the opening of the foodcup. Orienting responses were video recorded and scored by an independent observer. Both responses were analyzed over a 15s interval: the 5s before the light CS illumination (preCS) and in two blocks of 5s during the CS illumination (CS1 and CS2). Typically, orienting scores are higher in CS1 and foodcup behavior is higher CS2 (Hilz et al., 2019; Olshavsky et al., 2014). Therefore, orienting responses from CS1 and foodcup behavior from CS2 were analyzed; these scores were adjusted for baseline responding by subtracting preCS responses CS1 and CS2, respectively.

### Histology

Immediately following the final behavioral test rats were anesthetized with a 0.25ml pentobarbital and phenytoin solution (MED-PHARMEX Inc., Pomona, CA) intraperitoneal injection and transcardially perfused with 0.9% saline followed by 4% paraformaldehyde in 0.1 M phosphate buffer. Brains rested for 24 hours at 4°C in paraformaldehyde with 20% sucrose solution prior to rapid freezing with dry ice and storage at −80°C. Brains were sliced into 35µm coronal sections using a microtome and collected into 6 adjacent series; each series was mounted on Superfrost Plus microscope slides (Fisher Scientific, Pittsburg, PA) and stored at −80°C in sealed 3×1 inch microscope slide boxes (Fischer Scientific).

### Fluorescence *in situ* Hybridization (FISH)

All FISH procedures occurred in an RNAse free environment using procedures modified from Lee et al., 2016. One full brain series from each rat containing the mPFC, amygdala, dHPC, PVT, and vHPC was processed; two brains were excluded due to tissue damage. The plasmid used for generating the Arc antisense riboprobe contained the full-length cDNA (∼3.0 kbp) of *Arc* transcript. DNA was cut with 10x digestion buffer (NEBuffer; Biolabs; Ipswich, MA, USA) and 10x EcoRI restriction enzyme (Biolabs) in nuclease free water (Ambion). Following purification in EtOH, the DNA pellet was centrifuged, washed in EtOH, and resuspended in a TE buffer. The cRNA probe was made using T7 RNA polymerase (Ambion; Grand Island, NY, USA) and digoxigenin-UTP (DIG RNA labeling mix; Roche Applied Science, Indianapolis, IN, USA). The riboprobe was purified with EtOH precipitation and mini Quick Spin Columns (Roche) then stored at −80°C.

Slides were submerged in 4% paraformaldehyde in 0.1 M phosphate buffer for 1 hour to encourage tissue stability. After rinsing in PBS, slides were pretreated with proteinase K and were dehydrated through a series of ascending ethanol dips ranging from 50% to 100% EtOH. Tissue sections were air-dried and coated with ∼300µl hybridization solution containing the cRNA probe. Slides were temporarily coverslipped and incubated with hybridization solution for ∼20 hours at 60°C. After hybridization, coverslips were removed and slides were gently washed in 4X SSC at 60°C before being treated with RNase at 30°C and then washed in descending concentrations of SSC ranging from 4X to 0.1X at 60°C. Slides underwent immunocytochemical processing in a humid chamber using the PerkinElmer TSA Fluorescein system (NEL701001KT; PerkinElmer, Waltham, MA, USA). Tissues were coated with blocking buffer for 30 minutes prior to anti-DIG-HRP conjugate for 2 hours. Slides were gentle washed and then coated with fluroscein tyramide reagent (FITC) for 30 minutes in a dark humid chamber. Finally, each slide was coverslipped using a mounting medium that contained the nuclear stain 4’,6-diamidino-2-phenylindole (DAPI) (Vectashield; Vector Lab, Burlingame, CA, USA) and stored at −20°C until image acquisition occurred.

### Image Acquisition and Analysis

Images were acquired using an Axio Scope A1 microscope (Zeiss; Thornwood, NY, USA) from a subset of rats (*n* = 4 per group, run order ABBA). These animals were chosen for FISH analyses because appetitive behavior was higher in this run order (but not in ABAB). Regions of interest (i.e., the mPFC, nuclei of the amygdala, dPHC, PVT, and vHPC) were identified with nuclear DAPI staining under a 10x objective using Swanson’s Atlas (2004). Once identified, both FITC and DAPI images were taken under 40x objective and these images were compiled using a custom macro script with ImageJ (NIH, Bethesda, MD) software. Because the microscope used was not confocal, samples were collected from a single plane at various levels for each region of interest. Specifically, one sample for each coordinate was taken from two subregions of the mPFC: the prelimbic (PL; +4.20mm, +3.60mm, +3.20mm from Bregma) and infralimbic (IL; +3.20mm and +2.80mm) cortices. One sample was taken from each coordinate of four nuclei of the amygdala: the basolateral nucleus (BLA; -2.00mm; -2.45mm, -2.85mm, -3.25mm), the lateral nucleus (LA; -2.45mm, -2.85mm, -3.25mm), the medial portion of the central nucleus of the amygdala (CeA) (mCeA; -1.53mm and -1.78mm), and the lateral portion of the CeA (lCeA; - 1.78mm and -2.00mm). Two samples for each coordinate were taken from two regions of the dHPC: dCA1 and dCA3 (2.45mm, -2.85mm, -3.25mm). One sample for each coordinate was taken from two subregions of the PVT: the anterior PVT (i.e., PVTa; -1.08mm, -1.33mm, -1.53mm, - 1.78mm, -2.00mm) and posterior PVT (i.e., PVTp; -2.85mm, -3.25mm, -3.70mm). Two samples for each coordinate were taken from two subregions of the vHPC: vCA1 and vCA3 (-4.45mm, - 4.60mm, -5.00mm, -5.25mm).

Image analysis was conducted by a blind observer using the cell counter plugin with ImageJ software. First the total number of DAPI-stained cells present in an image were counted; next, DAPI-stained cells containing two clear *Arc* intranuclear foci were counted and classified as “nuclear” *Arc* expression. Then DAPI-stained cells containing diffuse perinuclear *Arc* staining were counted and classified as “cytoplasmic” *Arc* expression. Finally, DAPI-stained cells containing both nuclear and cytoplasmic *Arc* expression were counted and classified as “double” *Arc* expression. The number of double cells was then subtracted from the number of nuclear and cytoplasmic cells; this produced a final product of nuclear only, cytoplasmic only, and double only *Arc* expression; these *Arc*+ cells were expressed as a percentage of total DAPI-stained cells and then averaged across samples for subsequent statistical analyses.

### Statistical analyses

All statistical analyses were conducted in R (R Core Team, 2014).

#### Behavior

Appetitive conditioned foodcup and orienting behavior over conditioning, extinction, LTM and renewal testing was analyzed using repeated measures factorial ANOVA. Factors differed depending on the condition analyzed (see results), and effect size measured as partial eta-squared (η^2^_p_) was provided for significant ANOVAs. Bonferroni corrected *t-*tests were used to analyze *post hoc* comparisons on significant ANOVAs. A measure of effect size, Hedges’s *g* (for unequal sample size) or Cohen’s *d* (for equal sample size), was provided for significant *post hoc* tests for these and all subsequent analyses.

#### Arc mRNA expression

The percentage of DAPI cells expressing either cytoplasmic, nuclear, or co-expressing cytoplasmic and nuclear *Arc* mRNA was analyzed in each region of interest was analyzed using separate between-subjects’ factorial ANOVAs with the factor Group (i.e., Control, P-different, P-same, M/D-different, M/D-same); when a significant difference was detected *post hoc* comparisons were analyzed using Tukey’s HSD. The relationship between percentage of *Arc* mRNA expression and conditioned foodcup behavior at test (i.e., LTM or Renewal Z-score) was also examined using linear regression analysis. The factors Cycle and Status were compressed into Group to appropriately compare experimental rats to control rats that did not have similar cycle-status designations.

#### Context representation

To determine which, if any, regions might respond more or less to the different context exposures repeated measures ANOVA was used to compare Expression Type (i.e., cytoplasmic, nuclear, or co-expression) between experimental groups (i.e., using Cycle and Status as factors). Control rats did not have separate context exposure and therefore excluded from this analysis. Bonferroni corrected *t-*tests were used to analyze *post hoc* comparisons on significant ANOVAs.

## Results

### Conditioned Responding

#### Appetitive conditioning

The 16 daily appetitive conditioning trials were analyzed in blocks of 8 averaged trials with the exclusion of the unpaired block 1 (when the light CS was presented without the food US) for conditioned foodcup and conditioned orienting behavior. Factors included Training Blocks (i.e., blocks 2-8), Cycle (i.e., P or M/D), Status (i.e., Same or Different), and Test Order (i.e., ABBA or ABAB). *Conditioned orienting.* A significant main effect of Training Blocks indicated all groups acquired conditioned orienting behavior (*F*(6,180) = 7.00, *p* < .001, η^2^_p_ = .19; Fig. 2 Top Left). A significant interaction between Cycle, Status, Block, and Test Order (*F*(6,180) = 2.64, *p* = .01, η^2^_p_ = .08) suggested conditioned orienting differed depending on experimental group and test order. To determine if groups differed at the end of conditioning, a separate 2×2×2 factorial ANOVA was conducted on the final acquisition block (i.e., block 8). No main effect of Cycle, Status, Test Order, or interactions between these variables (*F*(1,31) = 0.76, 1.80, 0.03, 0.75 respectively, all *p* > .1) indicated all groups had a similar level of conditioned orienting at the end of conditioning. *Conditioned foodcup.* A significant main effect of Training Blocks indicated all groups acquired food-cup behavior (*F*(6,180) = 19.96, *p* < .001, η^2^_p_ = .40); Fig. 2 Bottom Left).

**Fig. 2.**
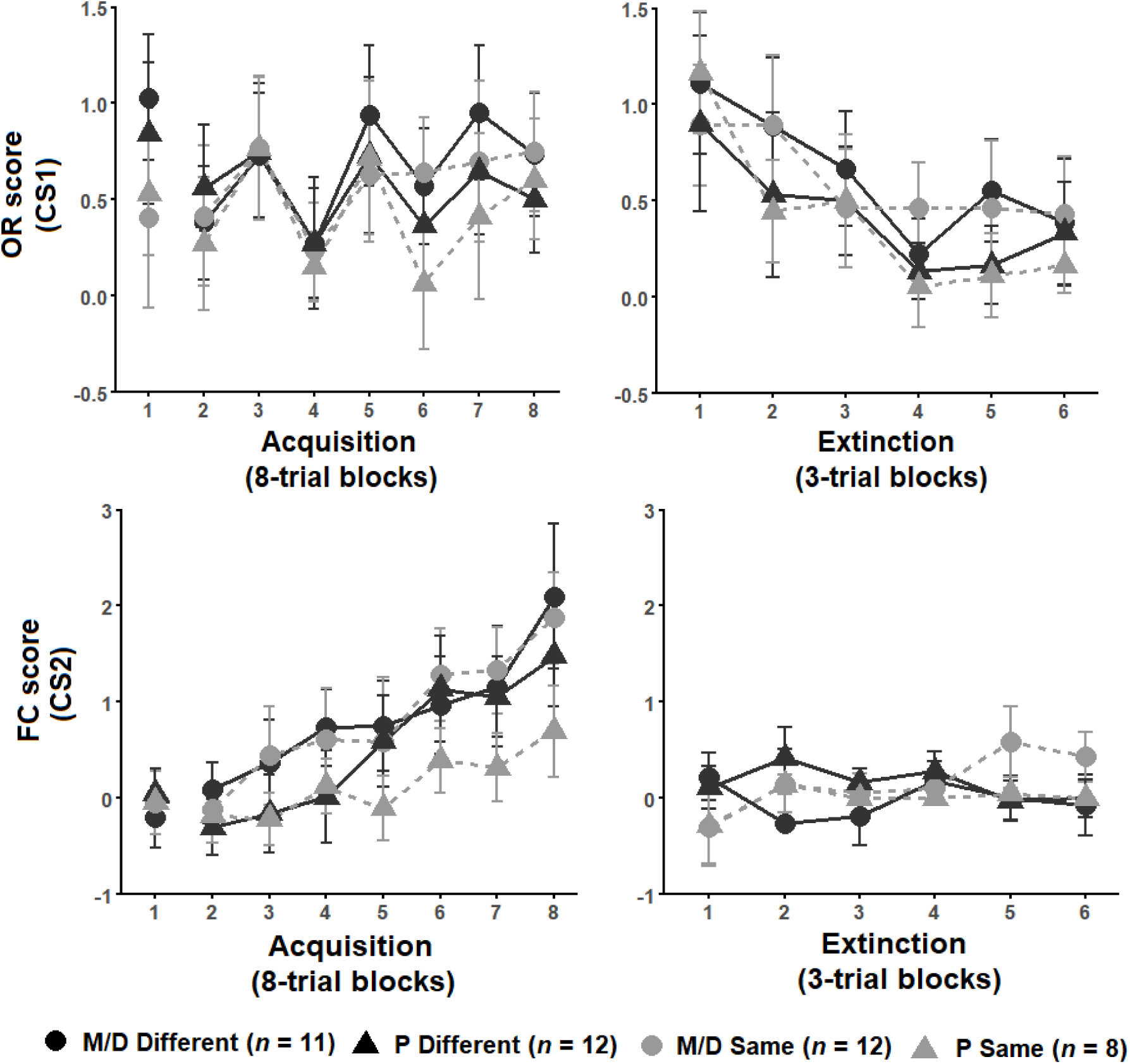
**(Top row).** Orienting responses (OR) +/- SEM for acquisition (left) and extinction training (right). Rats acquired ORs over acquisition and extinguished ORs over extinction training similarly between groups. **(Bottom row).** Foodcup (FC) score +/- SEM for acquisition (left) and extinction training (right). Rats significantly acquired FC responses and over acquisition blocks (2-8) but did not show a significant reduction in FC behavior over extinction training.

#### Extinction training

The 18 extinction training trials were analyzed in blocks of 3 averaged trials for conditioned foodcup approach and conditioned orienting. Factors included Extinction Blocks (i.e., blocks 1-6), Cycle, Status, and Test Order. *Conditioned orienting.* A significant main effect of Block (*F*(5,135) = 6.08, *p <* .001, η^2^_p_ = .20) but no interactions of Block with Cycle or Status indicated groups similarly decreased orienting responses over extinction blocks (Fig. 2 Top Right). *Conditioned foodcup.* Groups did not significantly change foodcup behavior over the extinction training as indicated by a nonsignificant main effect of Block (*F*(5,160) = 0.90; Fig. 2 Bottom Right).

#### Renewal

The 3 trials of renewal were compared to the 3 trials of the LTM probe and the last 3 trials of extinction for conditioned foodcup and conditioned orienting behavior. Factors included Condition (i.e., EXT, LTM or Renewal), Cycle, Status, and Test Order. *Conditioned Orienting.* A significant main effect of Condition (*F*(2,270) = 4.20, *p* = .01, η^2^_p_ = .03) and significant interactions between Condition and Test Order (*F*(2,270) = 2.95, *p* = .05, η^2^_p_ = .02) indicated differences in orienting responses depending on the order of tests. *Post hoc* analyses with an adjusted alpha of *p* < .008 indicated orienting responses at LTM were significantly higher than at EXT across all groups (*p* = .006, main effect); no other relationships were significant at the adjusted level (Fig. 3 Top Row). *Conditioned foodcup.* A significant main effect of Test Order (*F*(1,31) = 4.34, *p* < .05, η^2^_p_ = .12) was detected along with significant interactions between Cycle, Status, and Condition (*F*(2,303) = 3.05, *p* < .05, η^2^_p_ = .14) and Cycle, Condition, and Test Order (*F*(2,303) = 4.13, *p* = .01, η^2^_p_ = .18); a non-significant main effect of Conditioned was also detected (*F*(2,303) = 2.73, *p* = .07). *Post hoc* tests with an alpha level of *p* < .025 were used to compare conditioned food-cup behavior for test order. Food-cup behavior was significantly higher in the renewal test when the test order was ABBA compared to ABAB (*p* < .01, *g* = 0.59). Because renewal was higher in ABBA, *post hoc* tests with an alpha level of *p* < .0125 were used to compare LTM and renewal between experimental groups in the ABBA test order and showed that P- different rats exhibited significantly more renewal of food-cup behavior (*p* < .01, *d* = 0.84) at the renewal test compared to the LTM test (Fig. 3 Bottom Row).

**Fig. 3.**
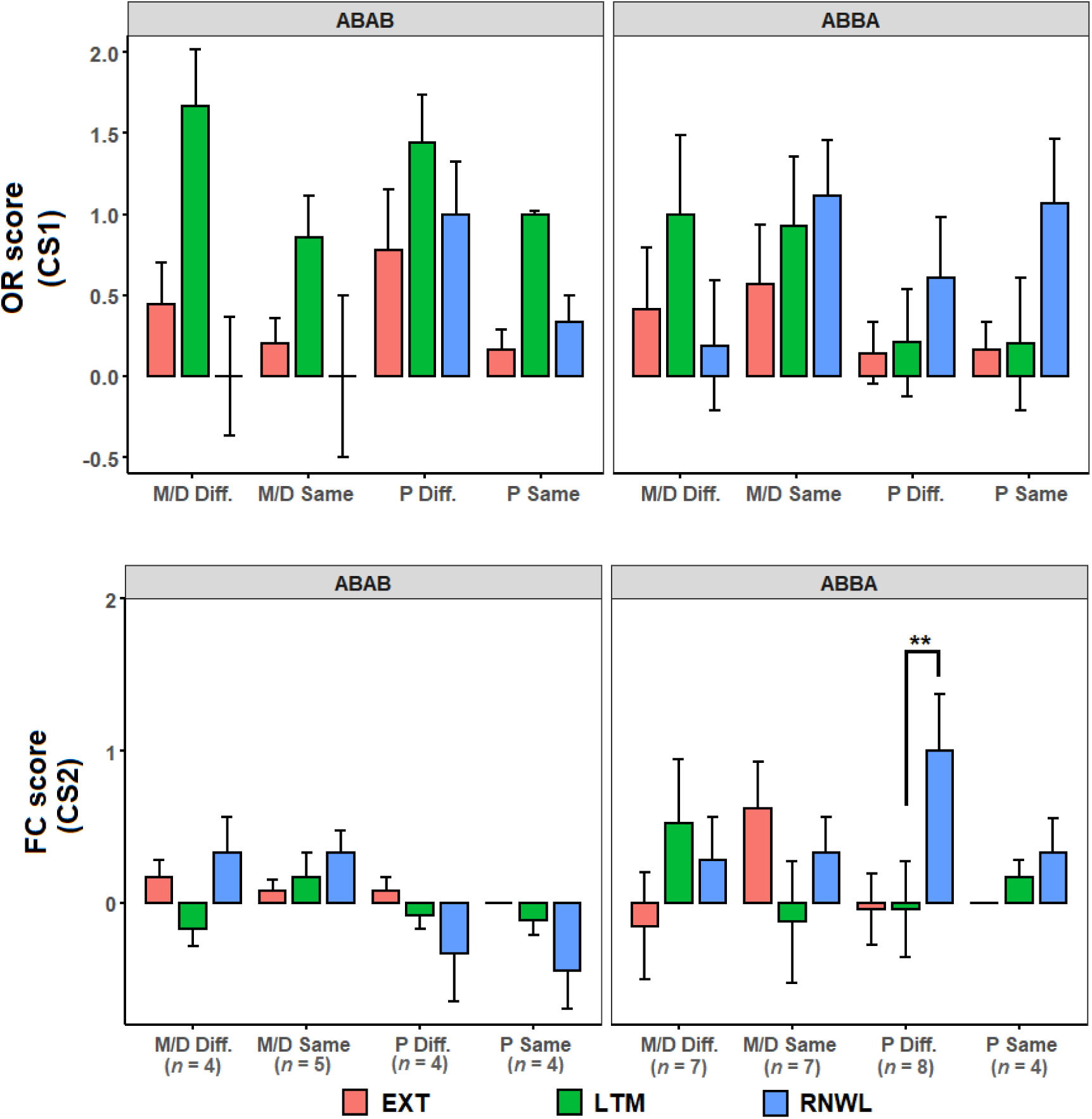
**(Top row).** OR score +/- SEM of last three extinction trials (EXT), LTM probe (LTM), and renewal test (RNWL) for ABAB test order (left) and ABBA test order (right). Rats showed more OR at LTM compared to EXT between groups. **(Bottom row).** FC score +/- SEM of last three extinction trials, LTM probe, and renewal test for ABAB test order (left) and ABBA test order (right). P-different rats exhibited significant return of FC behavior during RNWL compared to LTM (*p* < .01) in the ABBA test order.

#### *Arc* mRNA expression

For each region, the percent of DAPI-stained cells expressing nuclear, cytoplasmic, or co-expressing nuclear and cytoplasmic *Arc* mRNA (Fig. 4) were first compared between P and M/D home-cage controls using one-way ANOVA; no significant differences in *Arc* mRNA expression were observed based on the estrous cycle. Home-cage controls were combined into a single control group for subsequent analyses (*n* = 4).

**Fig. 4.**
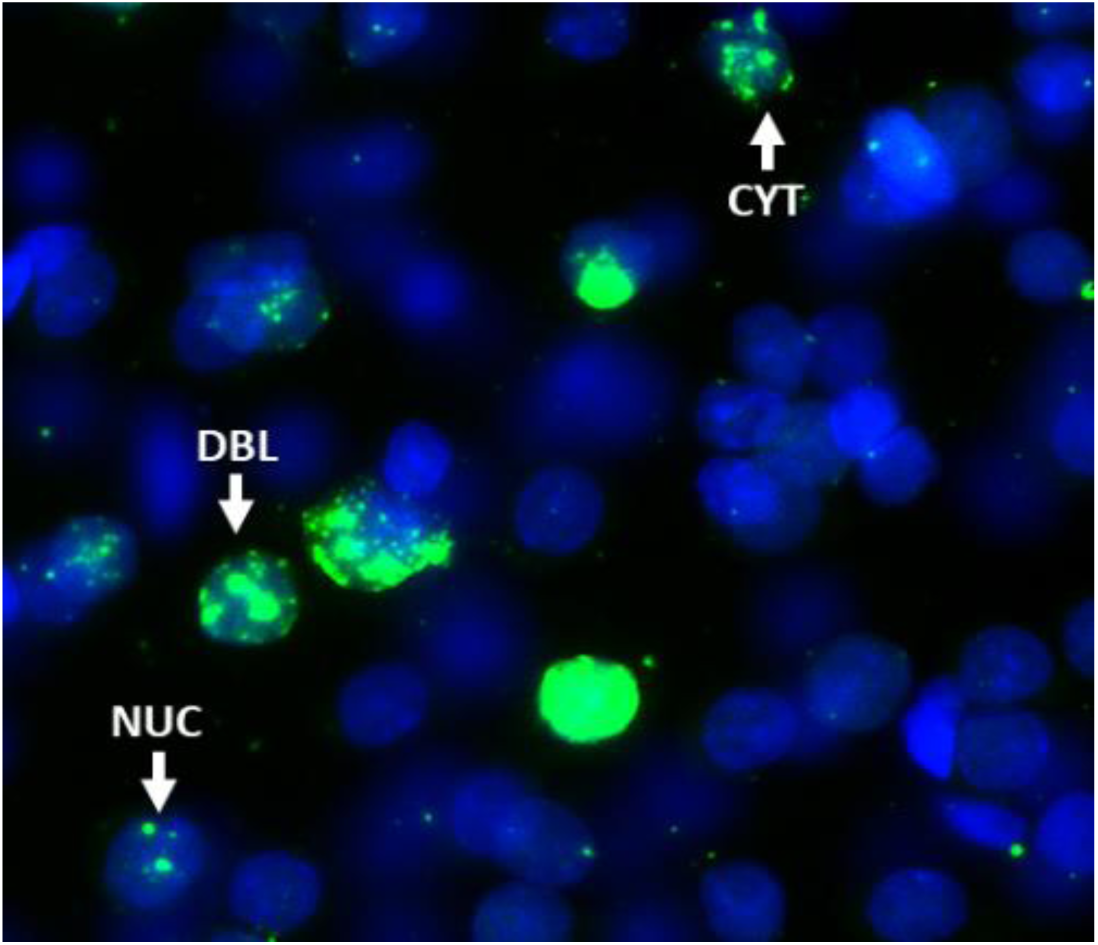
Example of tissue imaged after catFISH procedures showing DAPI stained cells (blue) from the PL region of mPFC. White arrows indicate cells expressing either nuclear (NUC), cytoplasmic (CYT), or co-expressing both nuclear and cytoplasmic (DBL) *Arc* mRNA (green). CYT corresponds temporally to the LTM context exposure, and NUC corresponds temporally to the renewal context exposure. DBL cells are responding to both context exposures.

#### Prefrontal Cortex

No differences were observed in the percentage of DAPI cells expressing either cytoplasmic or nuclear *Arc* mRNA between groups in the IL and PL cortices of the mPFC. A significant difference in the percentage of total DAPI cells co-expressing both nuclear and cytoplasmic *Arc* mRNA was detected between groups in both the PL (*F*(4,64) = 3.12, *p* < .05, η^2^_p_ = 0.16) and IL (*F*(4,60) = 3.48, *p* < .05, η^2^_p_ = 0.19). *Post hoc* analyses indicated no significant differences between groups in the PL but did suggest trends wherein M/D-same rats co-expressed a moderately higher percentage of *Arc* mRNA compared to M/D-different rats (*p* = .06) and moderately lower percentage of *Arc* mRNA compared to P-same rats (*p* = .07). Significant *post hoc* comparisons were observed in the IL, wherein P-different rats co-expressed a significantly higher percentage of *Arc* mRNA compared to both M/D-different (*p* < .05, *d* = 1.11) and P-same (*p* < .05, *d* = 1.22) rats; a trend was also observed that suggested P-different rats co-expressed a higher percentage of *Arc* mRNA compared to control (*p* = .08) rats (Fig. 5).

**Fig. 5.**
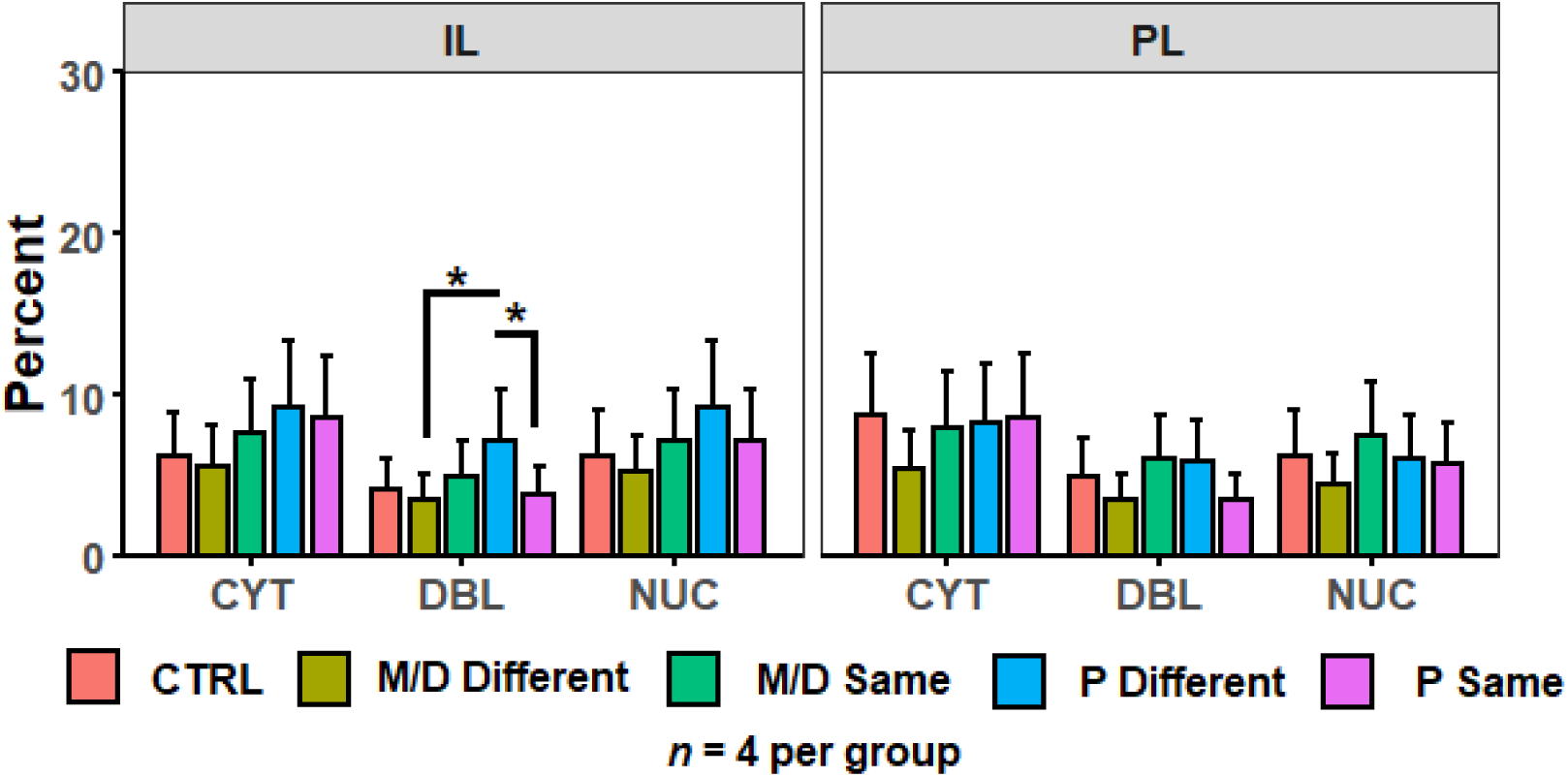
Percent of DAPI+ cells +/- SEM expressing cytoplasmic (CYT), double (DBL), or nuclear (NUC) *Arc* mRNA in the infralimbic (IL, left) and prelimbic (PL, right) cortices of the PFC. P-different rats had a significantly higher percentage of DBL cells in the IL.

#### Amygdala

No differences were observed in the percentage of DAPI cells expressing either cytoplasmic or nuclear *Arc* mRNA between groups in BLA, LA, and lCeA; in the mCeA the percentage of cytoplasmic (but not nuclear) *Arc* mRNA was significantly different between groups (*F*(4,81) = 2.65, *p* < .05, η^2^_p_ = .12). *Post hoc* analyses indicated control rats expressed a significantly higher percentage of cytoplasmic *Arc* mRNA compared to both M/D- and P- different rats (both *p* < .05, *d =* .91 and .95 respectively) and non-significantly compared M/D-same rats (*p* = .08). The percentage of total DAPI cells co-expressing both nuclear and cytoplasmic *Arc* mRNA did not differ between groups in the BLA (*F*(4,82) = 0.14) or mCeA (*F*(4,48) = 1.19). A non-significant trend suggested the percentage of total DAPI cells co-expressing nuclear and cytoplasmic *Arc* mRNA was moderately different between Groups (*F*(4,81) = 2.323, *p* = .06) in the lCeA. Significant differences in the percentage of total DAPI cells co-expressing nuclear and cytoplasmic *Arc* mRNA were observed between Groups in the LA (*F*(4,51) = 4.038, *p* < .01, η^2^_p_ = 0.24) and *post hoc* comparisons indicated the P-different rats co-expressed a significantly higher percentage of *Arc* mRNA compared to both control (*p* < .05, *d* = 1.36) and M/D-different (*p* < .01, *d* = 1.46) rats (Fig. 6).

**Fig. 6.**
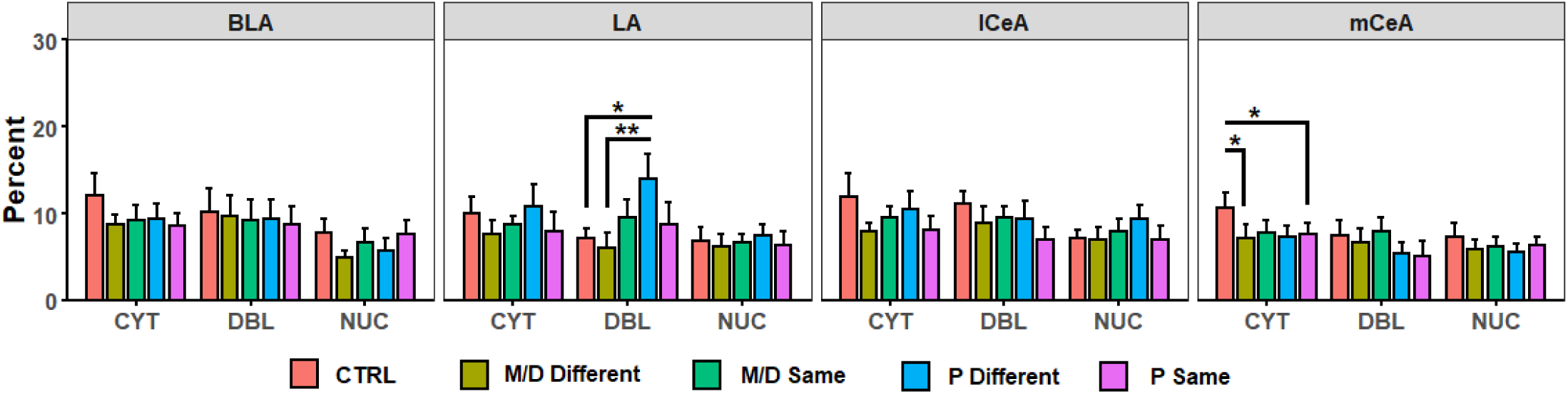
Percent of DAPI+ cells +/- SEM expressing CYT, DBL, or NUC *Arc* mRNA in the BLA, LA, lCeA, and mCeA regions of the amygdala. P-different rats had a significantly higher percentage of DBL cells in the LA compared to M/D-different and control (CTRL) rats. CTRL rats had a higher percentage of CYT cells in the mCeA compared to M/D-different and P-same rats in the mCeA.

The amygdala was the only region showing significant or moderate non-significant relationships between percentage of *Arc* mRNA expression and behavior (Fig. 7). The percentage of DAPI cells expressing nuclear *Arc* mRNA negatively predicted renewal Z-score in both the BLA (R^2^ = .31, β = -1.05 +/- 0.43, < .05; Fig. 7 Left) and mCeA (R^2^ = .22, β = -0.61 +/- 0.32, *p* = .07; Fig. 7 Right).

**Fig. 7.**
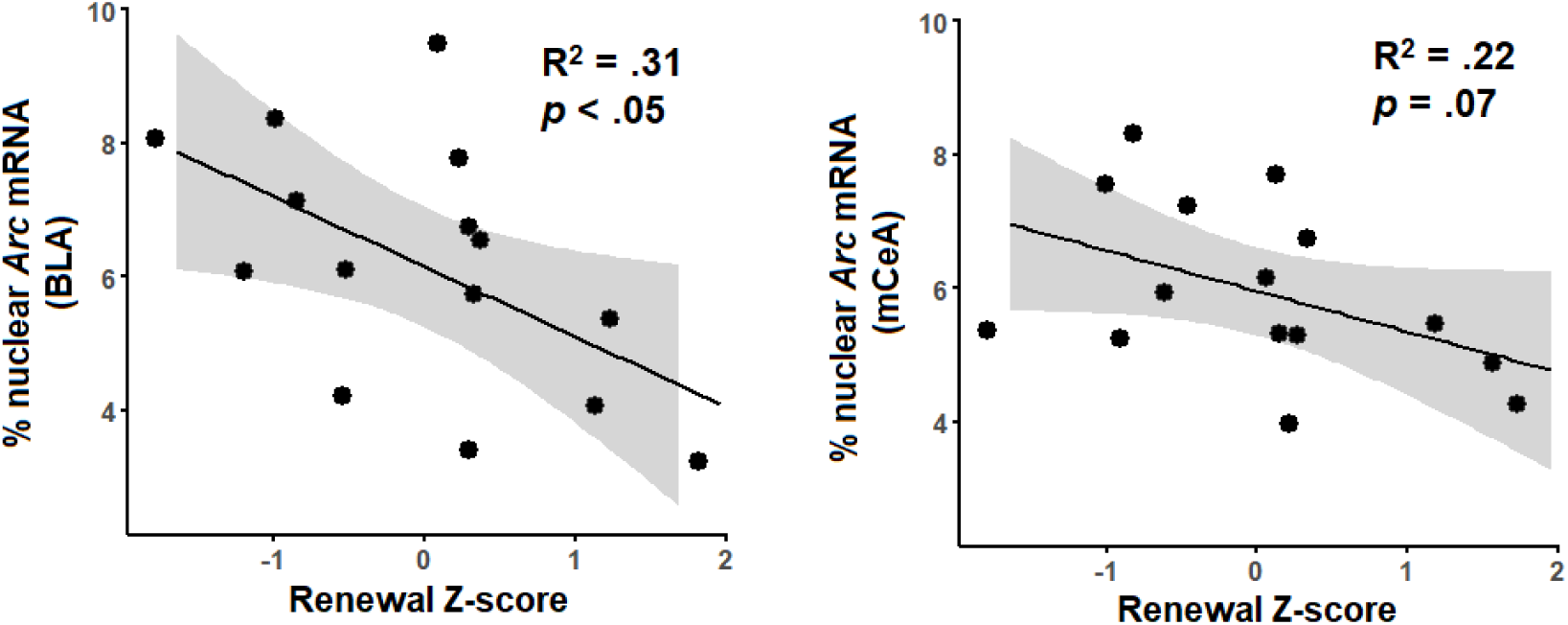
Relationship between the percentage of DAPI+ cells expressing nuclear *Arc* mRNA and normalized renewal Z-score in BLA (left) and non-significantly in mCeA (right). Regression line in black and standard error 95% confidence interval in gray.

#### Dorsal hippocampus

In dCA1 no differences were observed in the percentage of DAPI cells expressing cytoplasmic *Arc* mRNA between groups; however, the percentage of nuclear *Arc* mRNA non-significantly differed between groups (*F*(4,132) = 2.21, *p* = .07). In dCA3 there was a significant difference in the percentage of cytoplasmic *Arc* mRNA (*F*(4,126) = 3.14, *p* < .05, η^2^_p_ = .09) and a non-significant difference in percentage of nuclear *Arc* mRNA (*F*(4,126) = 2.20, *p* = .07). *Post hoc* analyses indicated P-different rats expressed a significantly higher percentage of cytoplasmic *Arc* mRNA compared to P-same (*p* < .05, *d* = .84) and non-significantly to control (*p* = .08) rats. A significant difference in the percentage of total DAPI cells co-expressing both nuclear and cytoplasmic *Arc* mRNA was detected between groups in both the dCA1 (*F*(4,132) = 5.17, *p* < .001, η^2^ = 0.14) and dCA3 (*F*(4,126) = 3.52, *p* < .01, η^2^ = 0.10). In dCA1, *post hoc* analyses indicated that M/D-same and P-same co-expressed a significantly lower percentage of *Arc* mRNA compared to control (*p* < .01, *d* = 0.78; *p* < .01, *d* = 0.77) rats; additionally, a trend was observed wherein M/D-different rats also co-expressed a lower percentage of *Arc* mRNA compared to control rats (*p* = .08). P-different rats co-expressed a significantly higher percentage of *Arc* mRNA compared to M/D-same rats (*p* < .05, *d* = 0.93) and a non-significantly higher percentage compared to P-same rats (*p* = .09). In dCA3, *post hoc* analyses indicated that P-same rats co-expressed a significantly lower percentage of *Arc* mRNA compared to both control (*p* < .05, *d* = 0.75) and P-different (*p* = .01, *d* = 1.40) rats (Fig. 8).

**Fig. 8.**
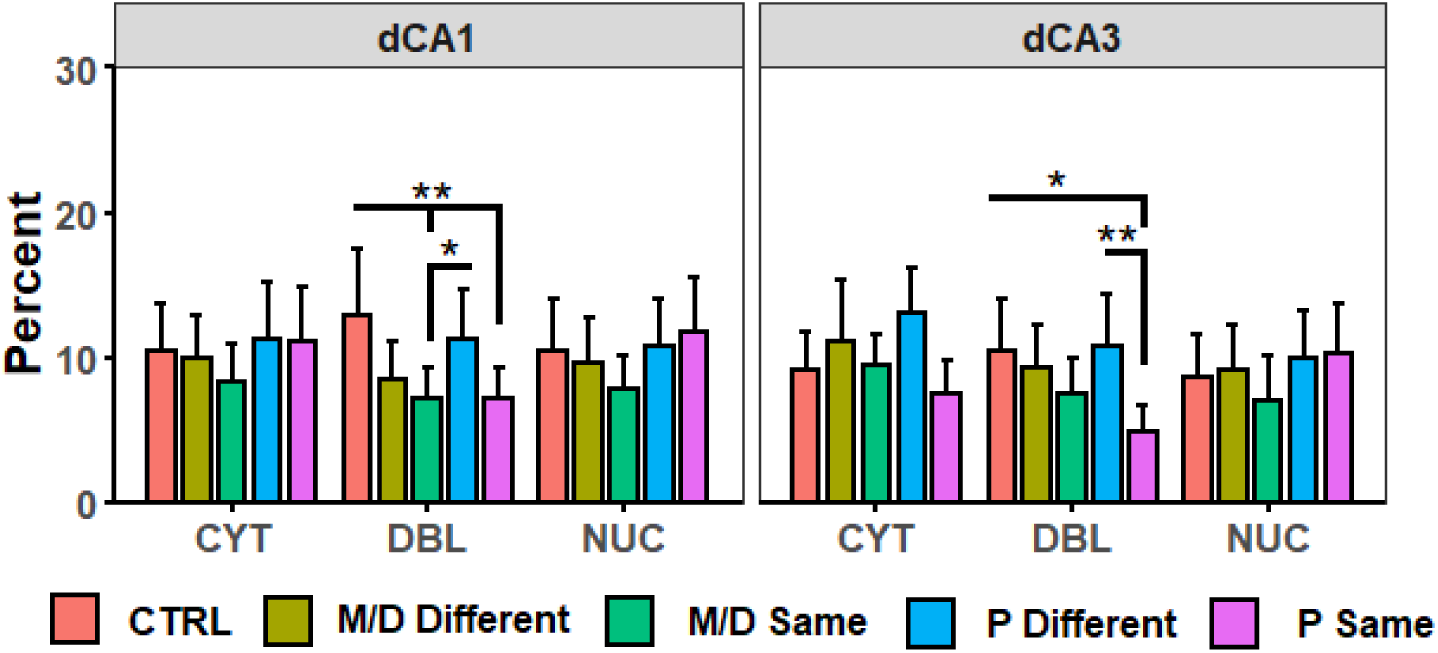
Percent of DAPI+ cells +/- SEM expressing CYT, DBL, or NUC *Arc* mRNA in dCA1 (left) and dCA3 (right) regions of the dHPC. In dCA1 CTRL rats had a higher percent of DBL cells than M/D and P-same rats; P-different rats also had a higher percent of DBL cells than M/D-same rats. In dCA3 P-same rats had a lower percent of DBL cells than CTRL and P-different rats.

#### Ventral hippocampus

No differences were observed in the percentage of DAPI cells expressing either cytoplasmic or nuclear *Arc* mRNA between groups in vCA1 and vCA3. The percentage of total DAPI cells co-expressing both nuclear and cytoplasmic *Arc* mRNA was significantly different between groups in the vCA1 (*F*(4,94) = 2.89, *p* < .05, η^2^_p_ = 0.11) but not the vCA3 (*F*(4,88) = 2.18, *p* = .07). *Post hoc* analyses indicated that P-same rats co-expressed a significantly higher percentage of *Arc* mRNA in the vCA1 compared to M/D-different (*p* < .01, *d* = 1.33) rats (Fig. 9).

**Fig. 9.**
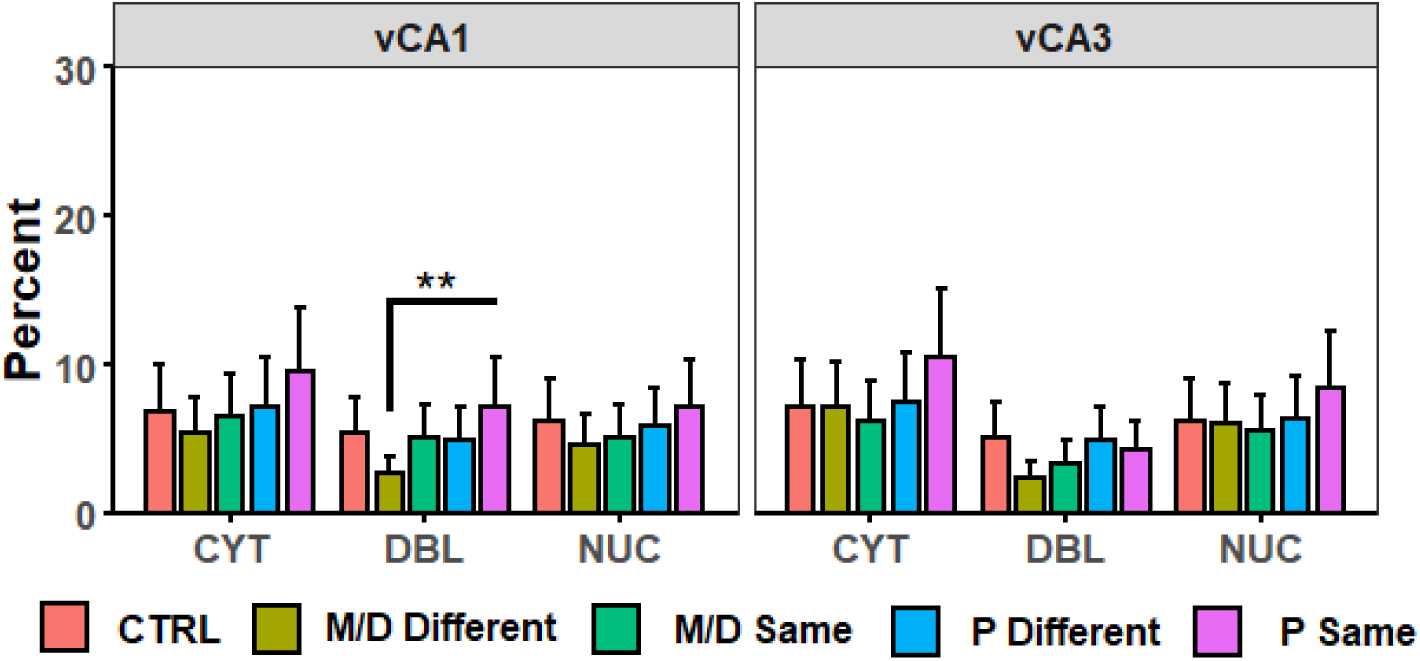
Percent of DAPI+ cells +/- SEM expressing CYT, DBL, or NUC Arc mRNA in vCA1 (left) and vCA3 (right) regions of the vHPC. P-same rats had a significantly higher percentage of DBL cells in the vCA1 compared to M/D- different rats.

#### Paraventricular nucleus of the thalamus

No differences were observed in the percentage of DAPI cells expressing either cytoplasmic or nuclear *Arc* mRNA between groups in PVTa and PVTp; moreover, the percentage of total DAPI cells co-expressing both nuclear and cytoplasmic Arc mRNA did not differ between groups in either PVTa nor PVTp (Fig. 10).

**Fig. 10.**
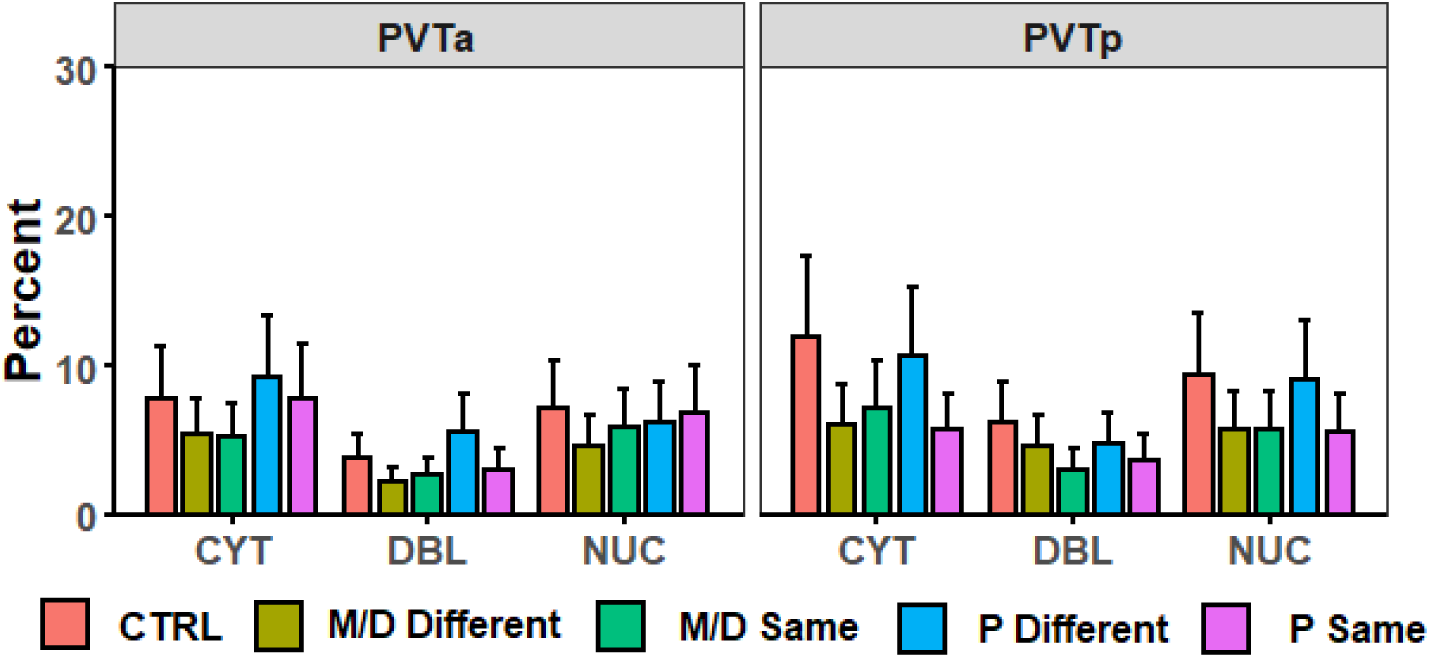
Percent of DAPI+ cells +/- SEM expressing CYT, DBL, or NUC *Arc* mRNA in PVTa (left) and PVTp (right). *Arc* mRNA expression did not differ between groups in either region.

#### Context representation

The results of the repeated measures analyses had a similar pattern wherein most regions (i.e., PL, IL, mCeA, LA, BLA, dCA3, vCA1, vCA3, PVTa, and PVTp) exhibited significant main effects of Expression Type (statistics detailed in Table 1). Additionally, dCA3 and vCA3 exhibited significant interactions between Cycle, Status, and Expression Type. *Post hoc* analyses with an alpha adjusted to 0.016 for main effects, 0.004 for within-subjects’ interaction terms, and 0.0027 for between-subjects’ interaction terms were conducted to determine the directionality of these effects. Generally, each region with a significant main effect of Expression Type had a higher percentage of DAPI cells expressing cytoplasmic *Arc* mRNA compared to nuclear and/or co-expression of *Arc* mRNA (main effect). Additionally, in the dCA3 P-same rats had a higher percent of nuclear and cytoplasmic *Arc* mRNA expression than co-expression and in vCA3 all experimental groups had a similar relationship of higher cytoplasmic than co-expression; M/D-different and P-same rats also showed more nuclear expression than co-expression in this region (within subjects). Finally, in dCA3 M/D-different rats had more nuclear expression of *Arc* mRNA compared to M/D-same rats, and P-different rats had more co-expression than P-same rats (between subjects).

**Table 1.** Within-subjects’ comparison of Arc mRNA expression patterns.

| Region | Comparison | $F(2,22)$ | $p$ | $\eta^2_p$ | *post hoc* |
| --- | --- | --- | --- | --- | --- |
| *Medial prefrontal cortex* |  |  |  |  |  |
| PL | Expression Type | 11.42 | < .001*** | 0.51 | CYT > NUC, $p = .003$<br>CYT > DBL, $p < .001$ |
| IL | Expression Type | 7.99 | < .01** | 0.42 | CYT > DBL, $p < .001$<br>NUC > DBL, $p < .001$ |
| *Amygdaloid nuclei* |  |  |  |  |  |
| mCeA | Expression Type | 3.93 | < .05* | 0.26 | CYT > DBL, $p < .01$<br>CYT > NUC, $p < .001$ |
| lCeA | Expression Type | 2.09 | > .1 | --- | --- |
| LA | Expression Type | 7.41 | < .01** | 0.4 | CYT > NUC, $p < .001$<br>DBL > NUC, $p < .001$ |
| BLA | Expression Type | 16.45 | < .001*** | 0.6 | CYT > NUC, $p < .001$<br>DBL > NUC, $p < .001$ |
| *Dorsal and ventral hippocampus* |  |  |  |  |  |
| dCA1 | Expression Type | 2.76 | 0.08 # | --- | --- |
| dCA3 | Expression Type | 8.35 | < .01** | 0.45 | CYT > DBL, $p < .001$<br>CYT > NUC, $p < .01$ |
| | Cycle*Status<br>*Expression Type | 4.52 | < .05* | 0.31 | P-same:<br>NUC > DBL, $p = .003$<br>P-same:<br>CYT > DBL, $p = .003$<br>M/D-different NUC > M/D-same NUC, $p < .001$<br>P-different DBL > P-same<br>DBL, $p = .001$ |
| vCA1 | Expression Type | 10.61 | < .001*** | 0.49 | CYT > DBL, $p < .001$<br>CYT > NUC, $p < .001$ |
| vCA3 | Expression Type | 32.94 | < .001*** | 0.75 | CYT > DBL, $p < .001$<br>CYT > NUC, $p < .001$<br>NUC > DBL, $p < .001$ |
| | Cycle*Status<br>*Expression Type | 4.48 | < .05* | 0.29 | M/D-same:<br>CYT > DBL, $p = .002$<br>M/D-different:<br>NUC > DBL, $p < .001$<br>M/D-different:<br>CYT > DBL, $p < .001$<br>P-different:<br>CYT > DBL, $p = .002$<br>P-same:<br>NUC > DBL, $p < .001$<br>P-same:<br>CYT > DBL, $p = .003$ |
| *Paraventricular nucleus of the thalamus* |  |  |  |  |  |
| PVTa | Expression Type | 23.5 | < .001*** | 0.68 | CYT > DBL, $p < .001$<br>NUC > DBL, $p < .001$ |
| PVTp | Expression Type | 10.89 | < .001*** | 0.5 | CYT > DBL, $p < .001$<br>NUC > DBL, $p < .01$ |

*Note.* Results list significant main or interaction effects of repeated measures ANOVA for each brain region, alpha ( $p$ ), effect size ( $\eta^2_p$ ), and *post hoc* results indicating directionality from Bonferroni corrected $t$ tests.

## Discussion

The results of this experiment replicate previous findings with some notable caveats. Estrous cycle stage (i.e., P or M/D) was considered during two important learning and behavioral expression windows: at extinction training and during LTM/renewal testing. Rats that underwent extinction training in P and LTM/renewal testing in some different stage of the estrous cycle (i.e., P-different rats) exhibited more renewal of appetitive behavior compared to all other experimental groups. Importantly, this effect was only observed depending on the order testing contexts were presented. Furthermore, this observation was restricted to conditioned foodcup, and not conditioned orienting, behavior. While P-different rats have been shown to renew conditioned foodcup (but not conditioned orienting) behavior in our pervious experiment (Hilz et al., 2019), the current experiment highlighted a previously unobserved test order effect in the renewal of appetitive behavior.

P-different rats only exhibited renewal when the order of test contexts was LTM followed by renewal (i.e., ABBA) as opposed to renewal followed by LTM (i.e., ABAB). In Hilz et al., 2019 no control for order of tests was included and all animals were conditioned and tested using an ABBA design; previous work in renewal of both appetitive and fear behavior that did control for potential test order effects suggested the order in which test contexts are presented would not influence behavioral expression (Trask and Bouton, 2016; Bouton and Schepers, 2015; Todd et al., 2012ab; Bouton et al., 2011; Rescorla 2008; Ji and Maren, 2005; Bouton and Ricker, 1994). The test order effect observed in this experiment is not in accordance with that body of research. The basis for the observed test order effect is difficult to determine, as groups did not differ in the acquisition of conditioned foodcup behavior nor was there any reason to believe the rapid test presentations would impair responding. It is possible that the small sample sizes associated with the ABAB counterbalance group drove the test order effect, and more systematic a robust exploration of counterbalanced renewal designs could provide illumination.

The results of this experiment extend previous findings that neuronal cell activity differs after renewal in a cycle-specific manner. In Hilz et al., 2019 P-different rats showed more cell activation (measured with FOS) in regions of the amygdala, PVT, and HPC after renewal testing. Here the immediate early gene *Arc* was examined, and differential *Arc* mRNA expression was observed in regions of the amygdala, dorsal HPC, and PFC. The pattern of *Arc* mRNA expression was similar to that of FOS in Hilz et al., 2019 with some notable differences: no differential FOS expression was observed in the PFC, but P-different rats did express more FOS in the ventral HPC. In this experiment differential expression and co-expression of *Arc* mRNA was observed in the IL region of the PFC, in the dorsal but not the ventral HPC, and in the amygdala. Nuclear *Arc* mRNA activity in the BLA and mCeA regions of the amygdala also predicted renewal of appetitive behavior. Moreover, it was observed that regions in this experiment were more responsive to the LTM context exposure than to the renewal context exposure, indicated by overall higher percentage of cytoplasmic than nuclear *Arc* mRNA expression.

The purpose of this experiment was to understand what may drive estrous-cycle specific differences in renewal both in terms of behavioral and subsequent cellular expression. There were two main hypotheses as to why P-different rats may exhibit renewal compared to other groups: 1) elevated sex steroid hormones estradiol and progesterone associated with P may enhance context information processing and encoding in P rats during extinction, and aid in subsequent context-differentiation to guide conditioned responding at testing; or 2) these elevated hormones around P may enhance extinction training in P rats during extinction, and recall of extinction training occurs in a state-dependent manner.

### Hippocampal context representation in renewal

High estradiol in P may facilitate encoding of the extinction context and aid in subsequent differentiation of the extinction context from the acquisition/renewal context via its actions on the HPC. Estradiol (and peaks of estradiol during the estrous cycle) modulate synaptic plasticity in the HPC (Kumar et al., 2015; Smith et al., 2009; Woolley, 2007); HPC activity is heavily implicated in the encoding and retrieval of context information (Neves et al., 2008; Ji and Maren, 2007; 2005; Antagnostaras et al., 1999; Philips and LeDoux, 1992). The effects of estradiol on HPC functional activity extend to HPC-dependent behavior; in particular, estradiol both endogenously and exogenously enhances spatial memory in females (Frick et al., 2018; Barha and Galea, 2010; Luine, 1998). The CA1 region of the HPC regulates both spatial memory and context information processing (Ji and Maren, 2008; Daumas, 2005; McHugh, 1996; Tsien et al., 1996), and infusions of estradiol into CA1 enhance spatial memory in female mice (Conrad et al., 2012; Li et al., 2004).

If rats in P during extinction encode information about the extinction context better than M/D rats, P rats would be better able to differentiate the extinction context from the acquisition context. Context is represented at a cellular level by separate cell populations in the HPC (Smith and Mizumori, 2006; Vazdarjanova and Guzowski, 2004; Wilson and McNaughton, 1993) and context representations act as a cue to guide conditioned responding in a context-appropriate manner (Fanselow, 1990; Balaz et al., 1982). Support for this hypothesis would be indicated by less overlap of nuclear and cytoplasmic *Arc* mRNA in the HPC of P rats: it can be inferred that less overlap indicates better or more specific context representation while more overlap would indicate the opposite. In this experiment P-different rats exhibited a higher percentage of active cell population overlap in both regions of the dorsal HPC – indicating that the separate LTM and renewal context exposures were inducing *Arc* mRNA expression in the same HPC cell populations. It is notable, however, that P-different rats also exhibited a higher specificity of cytoplasmic *Arc* mRNA in the dCA3. Cytoplasmic activity in these brains would temporally relate to the extinction/LTM context exposure; therefore, these rats may have had a better representation of the extinction context.

### Extinction learning and recall in renewal

The PFC, amygdala, and HPC contribute to extinction and renewal of reward-seeking in a variety of ways. Regions of the PFC are differentially active in renewal: IL is active in the formation and retrieval of extinction memory in both fear (Sierra-Mercado et al., 2011; Laurent and Westbrook, 2009) and appetitive (Mendoza et al., 2015; LaLumiere et al., 2010; Rhodes and Killcross, 2004) conditioning, while PL is implicated in the expression of associative fear (Sierra-Mercado et al., 2011; Corcoran and Quirk, 2007; Vidal-Gonzalez et al., 2006) and appetitive (Ashwell and Ito, 2014; Hayen et al., 2014) learning. The amygdala facilitates extinction of both rewarding and aversive learning (Schroeder and Packard, 2004); cell populations in the BLA and LA selectively activate in response to extinction-associated context information (Orsini et al, 2013) and this activity is context-specific after extinction (Hobin et al., 2003). The HPC, as discussed, is implicated in both fear and appetitive contextual information of processing, encoding, and context-dependent memory retrieval (Riaz et al., 2017; Hobin et al., 2006).

A growing body of evidence suggests that estradiol during extinction training, both via the estrous cycle and with exogenous treatment, enhances extinction recall (Graham and Daher, 2016; Graham and Milad, 2014; 2013; Zeidan et al., 2011; Milad et al, 2010; 2009; Chang et al., 2009). Injection of estradiol into the dorsal HPC prior to extinction training enhances contextual fear extinction; it has been supposed that estradiol acts on estrogen receptors in this region to enhance retrieval of context-specific extinction memory (Chang et al., 2009). Because internal states can act as contexts to cue renewal of extinguished behavior (Schepers and Bouton, 2017; Eich, 1995; Bouton et al., 1990; Ahlers and Richardson, 1985), it is possible that the estrous cycle hormonal state of the female rat may act as a context and guide subsequent extinction recall and/or conditioned responding. P is characterized by dramatic changes in sex steroid hormone level (Smith et al., 1975; Butcher et al., 1974) and likely separates itself internally from M/D. Rats that undergo extinction training while in P may associate extinction learning and the extinction context with the P stage of the estrous cycle: rats may recall extinction training and subsequently repress conditioned responding dependent on the similarity of the internal hormonal context at testing.

As *Arc* mRNA expression is more specific after context-dependent extinction learning in the amygdala and PFC (Orsini et al., 2013), support for this hypothesis would be indicated by P-different rats exhibiting less overlap of nuclear and cytoplasmic *Arc* mRNA in those regions. Again, an opposite result was observed: P-different rats exhibited more overlap or less specificity of cell population activity in the PFC (specifically in the IL cortex) and in the lateral amygdala (LA). Interestingly, nuclear expression of *Arc* mRNA (temporally correlated with exposure to the renewal context) in the BLA and non-significantly in the mCeA was a negative predictor of renewal of appetitive behavior. BLA specifically is a hub for gating information into the amygdala and mediates renewal of fear conditioning (Kim et al, 2017; Chen et al., 2017; Quirk et al., 2003). The negative relationship between BLA and mCeA nuclear *Arc* mRNA expression and renewal of appetitive behavior may indicate that activity in these regions is contributing to suppression of behavioral responding at the renewal test.

Ultimately, the results of this experiment did not fully elucidate the way hormonal states around the estrous cycle drives renewal in female rats. Behaviorally, P-different rats showed clear context differentiation and intact extinction learning indicated by suppressed conditioned responding in the extinction/LTM test context and a return of conditioned responding in the acquisition/renewal test context. Neuronally, however, the pattern of *Arc* mRNA expression did not support this differentiation of behavior depending on the context-stimulus exposures: P-different rats exhibited less specificity of cell activity in regions that support the renewal effect. One would expect that less specified *Arc* mRNA cell population activity would indicate poor context representation/differentiation in HPC and poor context-specific extinction memory representation/recall in the PFC and amygdala. This observed overlap of cell population activity may be a facet of differences in *Arc* mRNA recruitment compared to other immediate early genes (e.g., c-Fos). *Arc* mRNA is not induced by stimulus experiences in the same way that FOS is expressed (Kubik et al., 2007); therefore, the differences in *Arc* mRNA and FOS expression between this study and Hilz et al., 2019 indicate a difference in the induction of immediate early gene activity. *Arc* mRNA is necessary for neuronal LTP and LTD and is supposed to aid in the creation or strengthening of neuronal connections (Nikolaienko et al., 2018; Bramham et al., 2010); behaviorally, *Arc* is expressed differentially in response to context-dependent cue exposures and mediates memory formation and recall (Chia and Otto, 2013; Korb and Finkbeiner, 2011). Conceptualizing *Arc* as a marker of “new learning” or as necessary for the development of future behavior could alter the interpretation of this experiment: female rats exhibiting renewal may be creating or updating information surrounding the context learning and recall experience in overlapping cell populations throughout the renewal network.

There were some limitations in this experiment. Sample sizes, particularly in catFISH, were small (*n* = 4 per group) and limited to rats that underwent ABBA procedures. The observation of a test order effect limited our ability to interpret results. The catFISH technique allowed advanced temporal examination of cell activity; however, the image acquisition and quantification process was time-consuming and required substantial training, and the sensitivity of catFISH staining to background noise was high and required exclusion of multiple samples. Future research that seeks to understand the role of the estrous cycle in renewal of appetitive behavior may utilize alternative techniques to conceptualize context representation and extinction recall during renewal procedures: active recording from cell populations during context exposure and extinction training may elucidate the way cell populations are influenced by the estrous cycle to stimulate or suppress conditioned responding. Additionally, examination of other immediate early genes using catFISH may broaden our understanding of context representation or extinction memory recall as *Arc* mRNA is not necessarily best interpreted as simple marker of generalized neuronal activity.

